# Astrocytic lysosome deficits reduce alpha-synuclein degradation and induce spread of pathology

**DOI:** 10.1101/2025.08.29.673014

**Authors:** Lindsay M. Roth, Olga Morozova, Jan Stöhr, Jason Schapansky

## Abstract

Parkinson’s Disease (PD) is a neurodegenerative disorder caused by the loss of dopaminergic neurons in the substantia nigra due to Lewy body aggregates, primarily composed of misfolded alpha-synuclein (αSyn). While PD progression is thought to be driven by a prion-like spread of αSyn aggregates between neurons, the role of astrocytes remains unclear. Observations of αSyn pathology in PD patient astrocytes suggest their potential involvement in processing aggregates. To investigate this, we studied astrocytes’ interactions with αSyn pre-formed fibrils (PFFs) and their effects in astrocyte-neuron co-cultures on the spread of seed-competent αSyn. Primary astrocytes quickly internalized and degraded αSyn PFFs. However, degradation was significantly hindered by lysosome-compromising agents like chloroquine, Leupeptin, or CA-074. Adding astrocytes to neuron cultures reduced endogenous αSyn aggregation, indicating their role in mitigating αSyn pathology. When lysosome efficiency in astrocytes was compromised, their anti-seeding effect diminished. Moreover, lysosome-compromised astrocytes preloaded with αSyn PFFs enhanced αSyn pathology in neurons, whereas unimpaired astrocytes did not. These findings suggest astrocytes can modulate and contribute to αSyn pathology spread, playing a significant role in PD pathogenesis.

## Introduction

Synucleinopathies, such as Parkinson’s Disease (PD), Lewy body dementia (LBD), and multiple system atrophy, are a spectrum of neurodegenerative diseases defined by the shared pathological hallmark of intracellular accumulation and aggregation of alpha-synuclein (αSyn) in neurons and oligodendrocytes (Calabresi *et al*, 2023; Koga *et al*, 2021). In post-mortem PD and LBD brains, intraneuronal inclusions, termed Lewy bodies (LB) or Lewy neurites (LN), are rich in misfolded αSyn phosphorylated at serine 129 (pS129), as well as, lipid, lysosomal, and mitochondrial components (Ramalingam *et al*, 2023; Anderson *et al*, 2006; Fujiwara *et al*, 2002). It is believed that the misfolding of αSyn is the central pathogenic process in initiation and progression of PD. Once misfolded, αSyn aggregates are thought to propagate among connected brain regions in a prion-like manner, templating endogenous αSyn and leading to disease progression (Braak *et al*, 2003; Angot *et al*, 2010; Dehay *et al*, 2015; Jucker & Walker, 2018). PD etiology is still unclear but includes a complex involvement of genetic, environmental, and age-related factors (Pang *et al*, 2019) that potentially converge on αSyn pathogenesis. Furthermore, gene mutations in αSyn are often associated with both sporadic and familial PD such as the autosomal dominant G209A mutant that leads to overexpression of pathological A53T αSyn, producing more aggregate prone αSyn and thus increasing disease severity (Ohgita *et al*, 2022; Polymeropoulos *et al*, 1997). Multiple copies of the αSyn gene results in early pathogenesis, with disease onset directly correlating to αSyn expression (Stefanis, 2012). Lastly, PD patients in prodromal and clinical stages can be precisely diagnosed by the presence of aggregated, seeding competent αSyn in CSF and skin samples (Russo *et al*, 2021; Peng *et al*, 2023).

Under physiological conditions, αSyn is highly expressed in neurons and associates with neuronal membranes to regulate the recycling of synaptic vesicles (Bendor *et al*, 2013). In PD, accumulation of αSyn-rich LB and LN induce progressive loss of dopaminergic (DA) neurons projecting from the substantia nigra (Poewe *et al*, 2017), leading to gradual motor deficits such as tremors, muscle stiffness, and postural instability (Sveinbjornsdottir, 2016). PD research has primarily focused on this neuron-centric mechanism of DA loss; however, emerging research suggests that non-cell autonomous mechanisms play a significant role in PD etiology and disease progression (Panicker *et al*, 2021; Brash-Arias *et al*, 2024; Harackiewicz & Grembecka, 2024; Kam *et al*, 2020; Booth *et al*, 2017). A larger effort to understand the role of glia cells in PD pathogenesis might be a key to the development of disease-modifying therapeutics.

Astrocytes, the most abundant glia cell present in the brain, are critical for maintaining brain homeostasis and are highly involved in neuroinflammation, an emerging feature of PD pathogenesis (Araújo *et al*, 2022; Bartels & Leenders, 2007; Mena & Yébenes, 2008; Verkhratsky *et al*, 2017). Astrocytes have a robust endolysosomal system that is critical for release of gliotransmitters, modulating energy metabolism of astrocytes and neurons, pruning synapses, and clearing protein aggregates (Wong, 2020; Balta & Zunke, 2021). Even though αSyn is predominantly expressed by neurons, aggregated αSyn has been detected in astrocytes in PD brains, suggesting that αSyn secreted by neurons is taken up by astrocytes (Altay *et al*, 2022; Sorrentino *et al*, 2019; Braak *et al*, 2007; Wakabayashi *et al*, 2000; Hishikawa *et al*, 2001). Further, it suggests that astrocytes in PD are also impaired as they cannot clear these aggregates sufficiently leading to intracellular accumulation. Studies have shown that astrocytes can take up αSyn via the endolysosome pathway and localize to lysosomes, suggesting a role for astrocytes in the clearance of misfolded αSyn aggregates (Loria *et al*, 2017; Cavaliere *et al*, 2017; Filippini *et al*, 2019). Additionally, studies have reported that αSyn can be transferred from neurons to astrocytes *in vitro* which may help in the clearance of αSyn aggregates (Loria *et al*, 2017; Tsunemi *et al*, 2020; Domenico *et al*, 2019). In fact, many PD genome-wide association studies genes that encode proteins involved in lysosomal function, such as TMEM175, LRRK2, GCase, ATP13A2, Cathepsin D (Cat D), and Cathepsin B (Cat B), are highly enriched in astrocytes and their risk variants may alter critical astrocytic lysosomal functions in addition to neuronal impairments (Wang *et al*, 2021; Kim *et al*, 2023; Nalls *et al*, 2019; Booth *et al*, 2017). Recently, studies using induced pluripotent stem cells (iPSC)-derived astrocytes from PD patients harboring mutations in LRRK2, GCase, or ATP13A2 have shown alterations in clearance and metabolism pathways and inflammatory responses, suggesting a role for astrocytic lysosome dysfunction contributing to PD progression (Booth *et al*, 2019, 2017; Qiao *et al*, 2016; Sanyal *et al*, 2020; Tsunemi *et al*, 2020; Aflaki *et al*, 2020; Domenico *et al*, 2019; Jacquet *et al*, 2021; Pons-Espinal *et al*, 2024).

Here, we aimed to examine the consequences of lysosome deficiency in astrocytes on both the metabolism of αSyn pre-formed fibrils (PFFs) and the ramifications on neuronal αSyn pathology. Primary wild-type (WT) astrocytes, with little to no expression of endogenous αSyn, were used to track uptake and degradation of our PFFs templated from isolated PD brain samples. Primary neurons overexpressing A53T αSyn were utilized to identify endogenous pS129 αSyn aggregation induced by PFFs. These data demonstrate that αSyn PFFs are readily taken up and preferentially degraded by astrocytes, which prevents pS129 αSyn pathology in surrounding neurons. Disrupting lysosome function with a general insult, such as chloroquine (CQ), or with a more targeted approach, such as the irreversible Cat B inhibitor CA-074, results in accumulation αSyn PFFs in astrocytes. Subsequently, the ability of astrocytes to mitigate αSyn pathology in neurons is reduced and astrocytes can spread αSyn PFFs capable of seeding endogenous αSyn in neurons. This work demonstrates that lysosome-compromised (LC) astrocytes can contribute to the spread of αSyn pathology, and that enhancement of lysosomal function in astrocytes could help assuage progression of PD or other synucleinopathies.

## Materials and Methods

### Preparation of primary mouse CD-1 astroglia cell cultures

All experiments were performed in accordance with the guidelines set forth by AbbVie Animal Care and Use Committees. *A*strocytes were isolated from brains of postnatal day 1 CD-1 mice (Charles River Laboratories, Wilmington, MA) and plated in T75 flasks as previously described (Schildge *et al*, 2013). We purified the astrocytes using the “shake-off” method (McCarthy & Vellis, 1980). Three independent assays calculating the percentage of Vimentin+, Glial fibrillary acidic protein (GFAP)+, Aldehyde dehydrogenase 1 family member L1 (ALDH1L1)+, or Aquaporin-4 (AQP4)+ cells in culture averaged 93%, 80%, 73.5%, and 89%, respectively, suggesting most of the cells are astrocytes (Jurga *et al*, 2021). After purification, astrocytes were plated into 6 well plates or 96 well plates for immunoblotting or immunocytochemistry, respectively.

### Preparation of A53T αSyn transgenic mouse primary cortical neuron cultures

All experiments were performed in accordance with the guidelines set forth by AbbVie Animal Care and Use Committees. Mixed neuron and glia cultures were isolated from brains of embryonic day 16 A53T αSyn transgenic mice bred in-house. Brains were isolated and dissected cortices were incubated in 3 mL Hank’s Balanced Salt Solution with Neuronal Isolation Enzyme plus papain (Thermo Fisher Scientific, Waltham, MA) for 15 minutes in a shaking water bath at 37°C. Cells were then diluted to 10 mL total volume in plating media (Dulbecco’s Modified Eagle Medium, 10% fetal bovine serum and 1% penicillin streptomycin (P/S)) and centrifuged at 1500 rpm for 5 minutes. Aspiration, resuspension, and centrifugation of the cells occurred twice more and then resuspended cells were strained with a 70 um filter and counted. For mixed cultures (neurons and astrocytes), cells were plated at 35,000 cells/well into 96-well plates in plating media for 4 hours. Media was then changed to culture media (Neurobasal plus, B27 plus supplement, 1% P/S, and 1% GlutaMAX supplement). The time of plating is considered day in vitro (DIV) 0.

### Glia depletion protocol for A53T αSyn transgenic mouse primary neuron cultures

Following the preparation of a neuron and astrocyte cell suspension as described above, cells were diluted to 20 mL culture media and counted. Mitotic inhibitor, 5-Fluoro-2’deoxyuridine (FUdR, 1 µM) was added and the FUdR-containing cell suspension was incubated in a metal bead bath at 37°C for 1 hour. Following incubation, cells were recounted and centrifuged at 300 x g for 5 minutes. Cells were resuspended in culture media and plated at 55,000 cells/well into 96-well plates.

### Preparation of primary human astrocytes

Fetal human cortical astrocytes (ScienCell Research Laboratories, Carlsbad, CA) were cultured *in vitro* in astrocyte media (Cat # 1801, ScienCell Research Laboratories, Carlsbad, CA) in T-75 flasks until they reached 90% confluency. Cells were then incubated with 0.25% trypsin at 37°C for 3 minutes to lift cells. Following ultracentrifugation at 300 x g for 5 minutes, cells were counted and plated at 10,000 cells/well into 96-well plates or 500,000 cells/well into 6-well plates for immunocytochemistry and immunblotting experiments, respectively.

### Preparation of αSyn recombinant PFF aggregates and drug treatments

Recombinant WT α-syn was expressed in *E. coli* strain BL21(DE3) and purified as previously described (Ghee *et al*, 2005). Disease-relevant PFFs were generated by incubating insoluble αSyn seeds purified from human PD brains for 7 days, as previously described (Marotta *et al*, 2021). Protein concentration was determined by UV-Vis Nanodrop measurement after incubation with 4M Guanidine HCL. PFFs were diluted to 0.1 mg/mL in Dulbecco’s PBS (DPBS) and sonicated for 1 min at 30% amplitude with 1 sec on, 1 sec off pulses using a QSonica 500 water bath sonicator. pHrodo-labeled PFFs were generated by following the manufacturers protocol (Cat #: P36011, ThermoFisher) with a labeling ratio of 3:1. The unreacted dye was removed from solution by centrifuging the fibril solution at 20,000xg for 20 min, removing the supernatant and resuspending the pellet in PBS. The following drugs were used in this study: chloroquine (CQ, 25 µM, Millipore Sigma Cat #: C6628), leupeptin (Leu, 10 µM, Selleckchem Cat #: S7380), CA-074 (10 µM, Tocris Biosciences Cat #: 4863), or pepstatin A (Pep A, 10 µM, Tocris Biosciences Cat #: 1190). Cells were treated with αSyn PFFs for the time and concentration specified for each experiment.

### Immunofluorescence

All cell culture experiments for immunofluorescence were performed in 96-well plates. All experiments involving primary neurons were fixed and immunostained on DIV 21. Cultures were fixed with ice cold methanol for 10 minutes, rinsed with PBS, and incubated with 5% goat serum, 0.2% *T*riton-x100 in PBS to permeabilize and block cells for 1 hour. Then cells were incubated at 4°C degree overnight with primary antibodies. Antibodies used per experiment are specified in figure legends. A list of all primary antibodies used are as follows: microtubule-associated protein 2 (MAP2, Abcam, Cat #: ab5392, 1:3000); GFAP (Thermo Fisher Scientific, Cat #: 14-9892-82, 1:3000); pS129 (Cell Signaling Technologies, Cat #: 23706, 1:2000); Lysosomal associated membrane protein 1 (LAMP1) for mouse astrocytes (Biolegend, Cat #: 121602 1:200); LAMP1 for human astrocytes (Cell Signaling Technologies, Cat #: 15665, 1:200); total α-synuclein (MJFR1, Abcam, Cat #: ab138501, 1:2000) and Galectin-3 (Abcam, Cat #: A3A12, 1:200). The next day, cells were rinsed twice with PBS and incubated with secondary antibodies (Alexa Fluor 488, 568 or 647 [ThermoFisher] at 1:500 dilution) for 1 hour at room temperature. Following a 2x PBS rinse, cells were incubated with 1:2000 dilution of Hoechst in PBS for 10 minutes to stain nuclei. Cells are left in PBS for imaging to avoid drying out. Cells were imaged using an Opera Phenix high content confocal microscope (*Revvity*, Shelton, CT) affixed with 4 excitation lasers (405, 488, 561 and 640 nm) and emission filters (435-550, 435-480, 500-550, 570-630 and 650-760 nm). Images were captured at 20x or 40x magnification with at least 9 fields acquired per well. Quantification of marker areas or cells positive for stained markers was performed with algorithms made on the *H*armony analysis software (*Revvity*).

### Immunoblotting

Primary mouse astrocytes were grown in 6-well plate tissue culture dishes for immunoblot until they reached approximately 80% confluency. Whole cell extracts of primary mouse astrocyte cultures were prepared with ice cold RIPA buffer (ThermoFisher Scientific, Waltham, MA), containing protease and phosphatase inhibitor cocktails (Roche Diagnostics) followed by probe sonication. Protein concentrations were determined by bicinchoninic acid (BCA) assay (Thermo Fisher Scientific, Waltham, MA). Protein (15 ug/lane) was loaded onto a 12% Bis-Tris gel for separation. After separation, proteins were transferred to nitrocellulose membranes using an iBlot system (Thermo Fisher Scientific, Waltham, MA) and blocked in tris-buffered saline with Tween 20 (TBS-T) with 5% milk for 1 hour at room temperature. Membranes were incubated overnight at 4°C with primary antibodies in TBS-T with 5% milk. Primary antibodies to the following antigens were used: total αSyn (Syn1/clone 42, 1:500 dilution, BD Biosciences Cat #: 610787), total αSyn (MJFR1, 1:500 dilution, Abcam Cat #: ab138501), and β-actin (1:8,000 dilution, Millipore Sigma Cat #: A1978). Following 3x washes with TBS-T, membranes were incubated with corresponding antigen specific horseradish peroxidase-conjugated secondary antibodies (1:10,000 dilution) in TBS-T with 5% milk. Membranes were incubated with chemiluminescent substrate SuperSignal West Dura (ThermoFisher Scientific, Waltham, MA) for 5 minutes and then blots were visualized using a ChemiDoc MP Imaging System (Biorad, Hercules, CA). Densitometric analysis of band intensities were conducted using Fiji (NIH RRID: SCR_002285). All bands were normalized to the loading control as specified in each experiment.

### Statistical analysis

Data were analyzed using GraphPad Prism statistical software (version 10.0) and were graphically presented as a fold change from the average value of the normalizing group per experiment ± SD. The average value of the normalizing group is represented as a dotted line. The normalizing group is specified in the figure legends per experiment. All immunoblots were normalized to a loading control, β-actin. Each experiment was performed at least 3 times on separate 96-well or 6-well plates.

## Results

### Primary astrocytes readily take up and preferentially degrade αSyn PFFs via the endolysosomal pathway

Due to the presence of αSyn pathology in astrocytes in PD patients (Wakabayashi *et al*, 2000; Altay *et al*, 2022), we hypothesized that astrocytes may play a significant role in metabolism of αSyn PFFs. Using WT astrocytes allowed for a model in which we could monitor metabolism of external αSyn aggregates since astrocytes express little to no endogenous αSyn (Braak *et al*, 2007; Mori *et al*, 2002). To measure rate of uptake, mouse astrocytes were treated with pHrodo-tagged recombinant αSyn monomer or PFFs and imaged live by fluorescence every 2 hours for 24 hours. The pHrodo tag allows for fluorescence only in acidic compartments such as endosomes or lysosomes, therefore only active uptake is observed. Representative images of αSyn monomer- or PFF-treated astrocytes showed increased pHrodo+ fluorescence over time (Figure 1A). The number of phRodo+ spots over time (Figure 1B) or at 24 hours (Figure 1C) was significantly increased in astrocytes treated with αSyn PFFs compared to monomer, implying that aggregates are more actively internalized compared to soluble αSyn.

**Figure 1.**
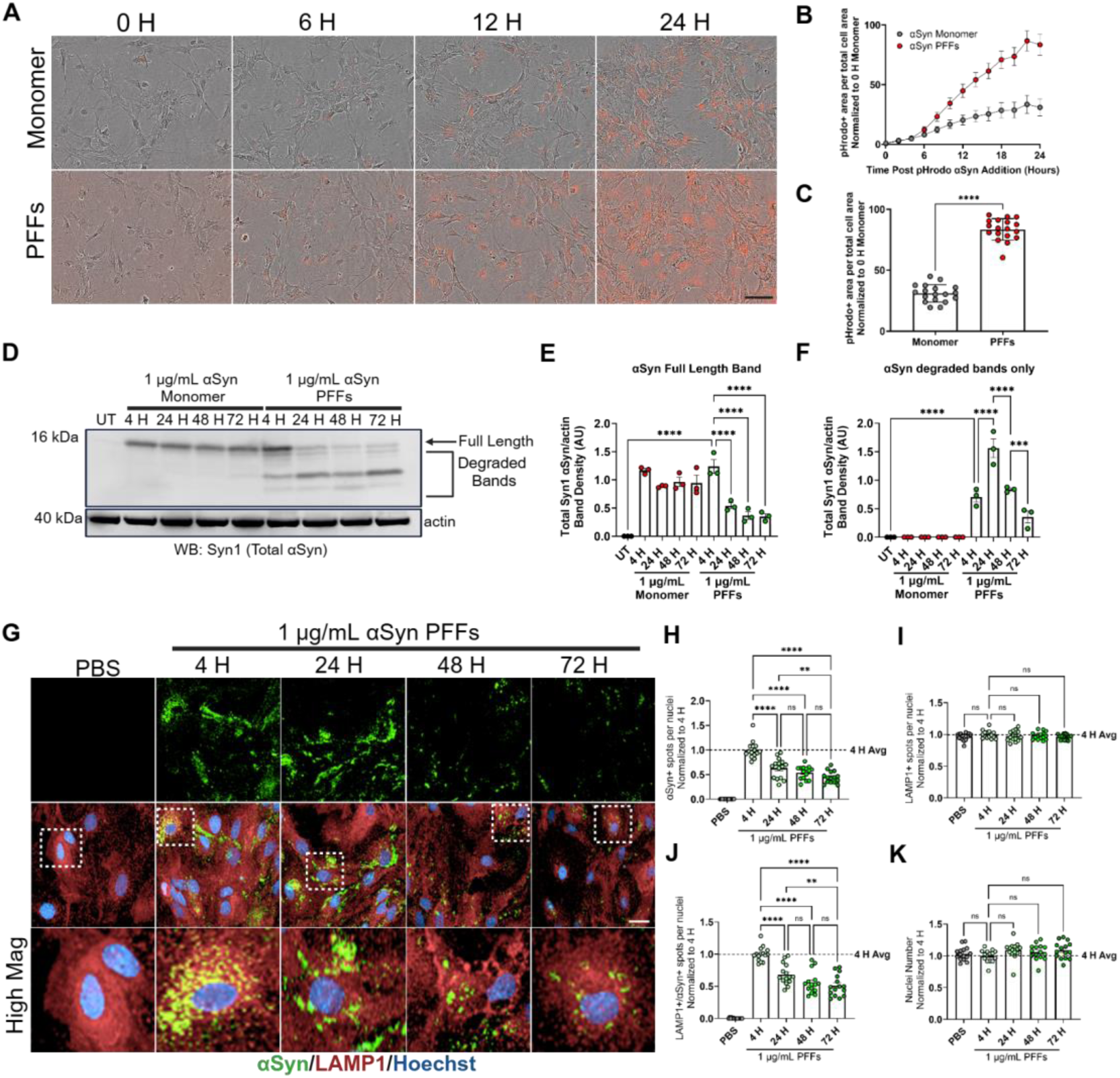
Primary astrocytes preferentiality take up and degrade αSyn PFFs via the lysosome. A) Bright-field images of astrocytes treated with phRodo-tagged αSyn monomer or PFFs overtime. Scale bar = 50 µm. Quantification of phRodo+ area per total cell area overtime (B) and at 24 H (C). Graphs B-C are expressed as a fold change of the 0 h monomer group. N = 6 wells/plate across 3 separate experiments. Unpaired student’s t-test, ****p = < 0.0001. D) Representative immunoblot of astrocyte lysates probed for total αSyn and β actin. Quantification of band intensities for full length total αSyn (E) and partially degraded αSyn (F). N = 3 separate experiments. One-way ANOVA with Sidak post hoc, ***p = < 0.001, ****p = < 0.0001. G) Representative images of immunocytochemically stained for total αSyn, LAMP1 and nuclei with Hoechst. Scale bar = 50 µm. Quantification of total αSyn+ spots per nuclei (H), LAMP1+ spots per nuclei (I), total αSyn+ and LAMP1+ spots per nuclei (J) and nuclei number (K). N = 3-6 wells/plate across 3 separate experiments. Graphs H-K are expressed as a fold change of the 4 H group. One-way ANOVA with Sidak post hoc, **p = < 0.01, ****= p < 0.0001.

Next, we assessed astrocytic degradation of αSyn monomer or PFFs by treating primary astrocytes with αSyn monomer or PFFs over time and immunoblotting for total αSyn. Total full length αSyn protein was initially present in both αSyn monomer- and PFF-treated astrocytes, but we only observed decreased αSyn abundance over time in αSyn PFF-treated astrocytes, indicating that aggregated, not monomeric, αSyn is targeted for degradation (Figure 1D, E).

Moreover, only αSyn PFF treatment resulted in the presence of degraded αSyn bands, represented by bands lower than 14 kDa, that increased after 24 h and then decreased at 48 and 72 h (Figure 1D,F). Using a C-terminal-localized αSyn antibody resulted in a loss of detection of degraded αSyn bands in the αSyn PFF-treated primary astrocyte lysates (Extended View Figure 1A,B), suggesting that astrocytes are degrading αSyn PFFs at the c-terminal end.

Given that αSyn PFFs were detected in acidic compartments, we asked whether we would observe decreased total αSyn abundance over time. Primary astrocytes were treated with αSyn PFFs over time and immunostained for total αSyn and the lysosome marker LAMP1 (Settembre *et al*, 2013). Subsequent quantification of αSyn PFF-treated astrocytes revealed fewer total αSyn+ spots over time and fewer total αSyn+ and LAMP1+ spots over time (Figure 1G, H, J), with no observed differences in total LAMP1+ spots or total nuclei over time (Figure 1I, J). These results are consistent with our immunoblotting results, suggesting an active digestion of αSyn aggregates that occurs quickly following exposure.

To confirm this was not a phenomenon localized to mouse astrocytes, primary human astrocytes were used to assess the differences in uptake and degradation of αSyn monomers or PFFs. Like mouse astrocytes, we observed that both monomers and PFFs are taken up, while only the PFFs were subsequently degraded (Extended View figure 1C-E). Additionally, the immunofluorescent assay results were consistent with the immunoblotting results, with decreased total αSyn+ spots and total αSyn+ and LAMP1+ spots in human primary astrocytes treated with αSyn PFFs (Extended View figure 1F-J). Thus, this was not a species-specific phenomenon but seems to relate to the human cells as well.

### αSyn PFFs accumulate in LC primary astrocytes

In figure 1 and extend view figure 1 we showed that mouse and human astrocytes take up and degrade αSyn PFFs via the endolysosomal pathway. We hypothesized that αSyn PFFs could accumulate if this pathway was perturbed. To examine this, we co-treated primary astrocytes with lysosome compromising agents and αSyn PFFs, followed by immunostaining for total αSyn after 24 hours. We utilized the lysosomotropic compound chloroquine (CQ, 25 µM), Cat D enzyme inhibitor pepstatin A (Pep A, 10 µM) or Cat B inhibitors leupeptin (Leu, 10 µM) and CA-074 (10 µM) to challenge lysosomal activity (Yoon *et al*, 2022; Gacko *et al*, 2007; Baici & Gyger-Marazzi, 1982; Gallagher *et al*, 2017). Quantification of challenged astrocytes revealed more total αSyn+ spots within lysosomes after 24 hours treatment compared to vehicle-treated astrocytes (Figure 2A,B). Interestingly, total αSyn+ spots in Pep A-treated astrocytes were similar to that of vehicle-treated astrocytes (Figure 2A, B), implying that aspartic acid proteases are not involved in astrocytic αSyn PFF degradation. Furthermore, astrocytes only demonstrated lysosome damage marker Galectin-3 (Gal-3) with CQ treatment as shown previously (Gallagher *et al*, 2017) and not with cathepsin inhibition (Extended View figure 2D-H), suggesting that lysosome damage is not required for aSyn accumulation. Moreover, the amount of αSyn remaining correlated with the severity of insult, with CQ having a greater effect. Therefore, both lysosomal damage and/or lysosomal enzymatic insufficiency can result in an accumulation of αSyn aggregates in astrocytes.

**Figure 2.**
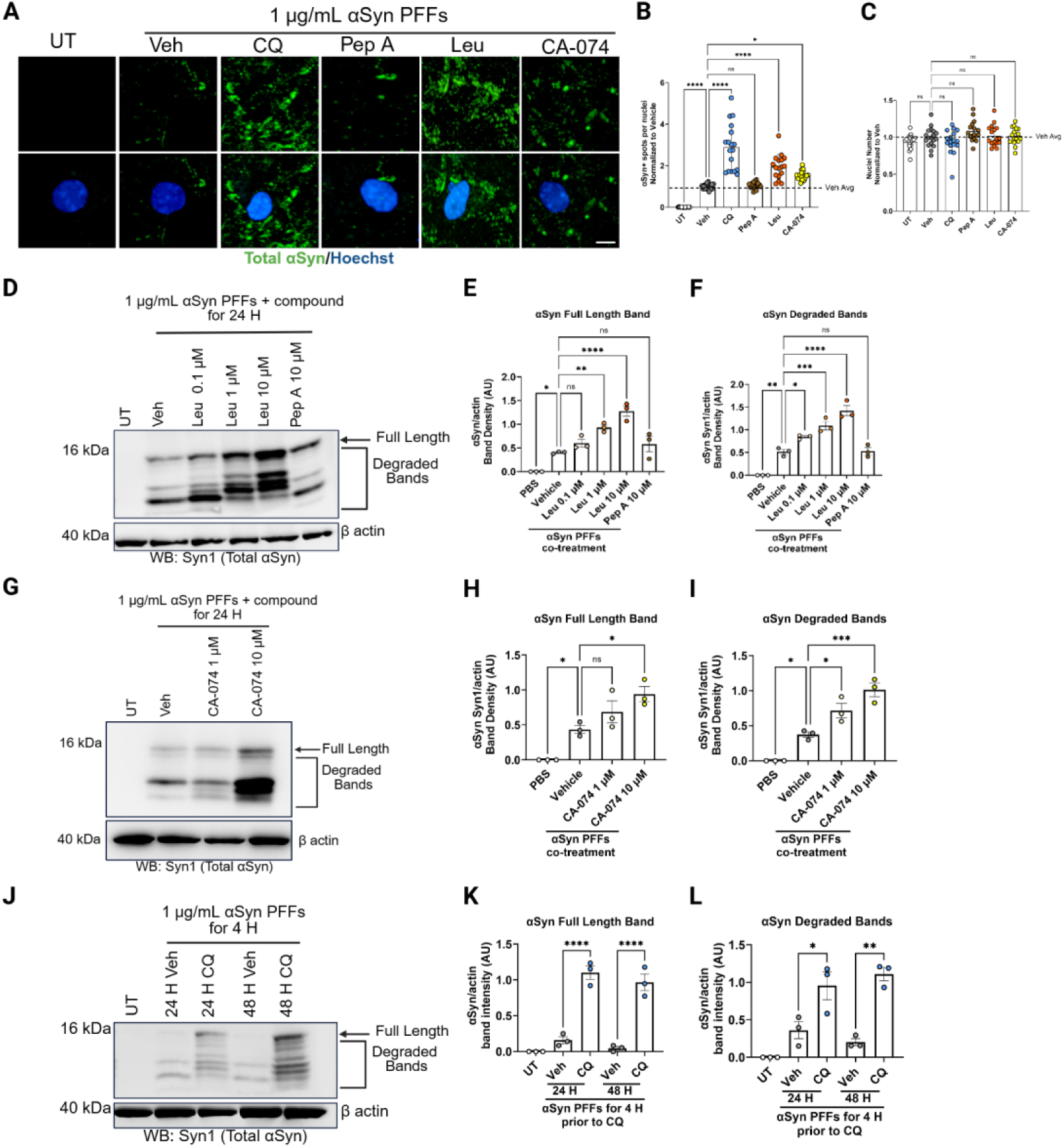
αSyn PFFs accumulate in lysosome-compromised (LC) primary astrocytes. A) Representative images of astrocytes immunocytochemically stained for total αSyn and nuclei with Hoechst. Scale bar = 50 µm. Quantification of total αSyn+ spots per nuclei (B) and nuclei number (C), expressed as a fold change of the vehicle. N = 3-6 wells/plate across 3 separate experiments. One-way ANOVA with Dunnett’s post hoc, *p = < 0.05, ****= p < 0.0001. D) Representative immunoblot of astrocyte lysates probed for total αSyn and β actin. Quantification of band intensities for full length non-degraded total αSyn (E) and partially degraded αSyn (F). N = 3 separate experiments. One-way ANOVA with Dunnett’s post hoc, *p = < 0.05, **p = < 0.01. G) Representative immunoblot of astrocyte lysates probed for total αSyn and β actin. Quantification of band intensities for non-degraded αSyn (H) and partially degraded αSyn (I). N = 3 separate experiments. One-way ANOVA with Dunnett’s post hoc, *p = < 0.05. J) Representative immunoblot of astrocyte lysates probed for total αSyn and β actin. Quantification of band intensities for non-degraded αSyn (K) and partially degraded αSyn (L). N = 3 separate experiments. One-way ANOVA with Sidak post hoc, *p = < 0.05, **p = < 0.01, ****p = < 0.0001.

To further elucidate the degradative capacity of lysosome-compromised (LC) astrocytes, we co-treated astrocytes with increasing concentrations of Leu (0.1, 1 or 10 µM), CA-074 (1 or 10 µM) or Pep A (10 µM) and αSyn PFFs for 24 hours and immunoblotted for total αSyn. Leu-treated astrocytes showed concentration-dependent accumulation of non-degraded αSyn compared to vehicle-treated astrocytes (Figure 2D,E). In addition to full length αSyn we also observed an accumulation of partially degraded αSyn suggesting that inhibition of Cat B reduces the efficiency of PFF degradation in the lysosome. The distinct banding patterns indicate a specific protease resistant confirmation of the αSyn fibrils (Figure 2D,F). In contrast we observed no difference in full length or degraded total αSyn in Pep A-treated astrocytes compared to vehicle-treated astrocytes suggesting that Cat D is not participating in the degradation of pathological αSyn (Figure 2D-F). Specific inhibition of cathepsin B with CA-074 also showed accumulation of non-degraded αSyn and distinct degraded αSyn bands compared to vehicle-treated astrocytes (Figure 2G-I). These results are consistent with the immunofluorescence results in which only Leu or CA-074-treated astrocytes increased abundance of total αSyn, suggesting that cysteine proteases and specifically cathepsin B function is critical for degrading αSyn aggregates in astrocytes.

Since CQ is a known inhibitor of endocytosis (Wang *et al*, 1993; Dutta & Donaldson, 2012), we also utilized a different paradigm to specifically demonstrate the effect of CQ on αSyn PFF degradation. In figure 2J, astrocytes were first preloaded with αSyn PFFs for 4 hours, rinsed and treated with vehicle or CQ for 24 or 48 hours. This data shows similar results to those in figure 2D-I, that both non-degraded αSyn and partially-degraded αSyn accumulated in CQ-treated astrocytes compared to vehicle-treated astrocytes (Figure K,L).

### Healthy astrocytes prevent pS129 pathology in neurons induced by αSyn PFFs

We have shown that astrocytes take up and preferentially degrade αSyn PFFs through the endolysosomal pathway, in a distinct manner from αSyn monomer. To model how this astrocytic ability may impact neuronal αSyn pathology, we developed multiple co-culture models. Figure 3A depicts the first co-culture assay we used to understand the effect of astrocytes on neuronal αSyn pathology. Briefly, DIV 7 A53T αSyn transgenic mouse neuroglia cultures were exposed to αSyn PFFs to induce endogenous αSyn aggregation. To ensure some uptake of αSyn PFFs into neurons, we waited four days after PFF treatment before adding increasing densities (5,000, 10,000, or 20,000) of mouse astrocytes to the neuroglia cultures. On DIV 21, the co-cultures were fixed and immunostained for astrocytes, neurons and endogenous αSyn pathology using GFAP, MAP2, or pS129, respectively. Representative images and quantification of astrocyte-added neuroglia cultures show fewer pS129+ spots compared with neuroglia cultures in the absence of additional mouse astrocytes (Figure 3B,C). Thus, the addition of exogenous astrocytes was protective, preventing neuronal-based αSyn aggregation. This assay includes A53T αSyn transgenic astrocytes, and adding exogenous astrocytes significantly decreases neuronal αSyn pS129+ pathology compared to control.

**Figure 3.**
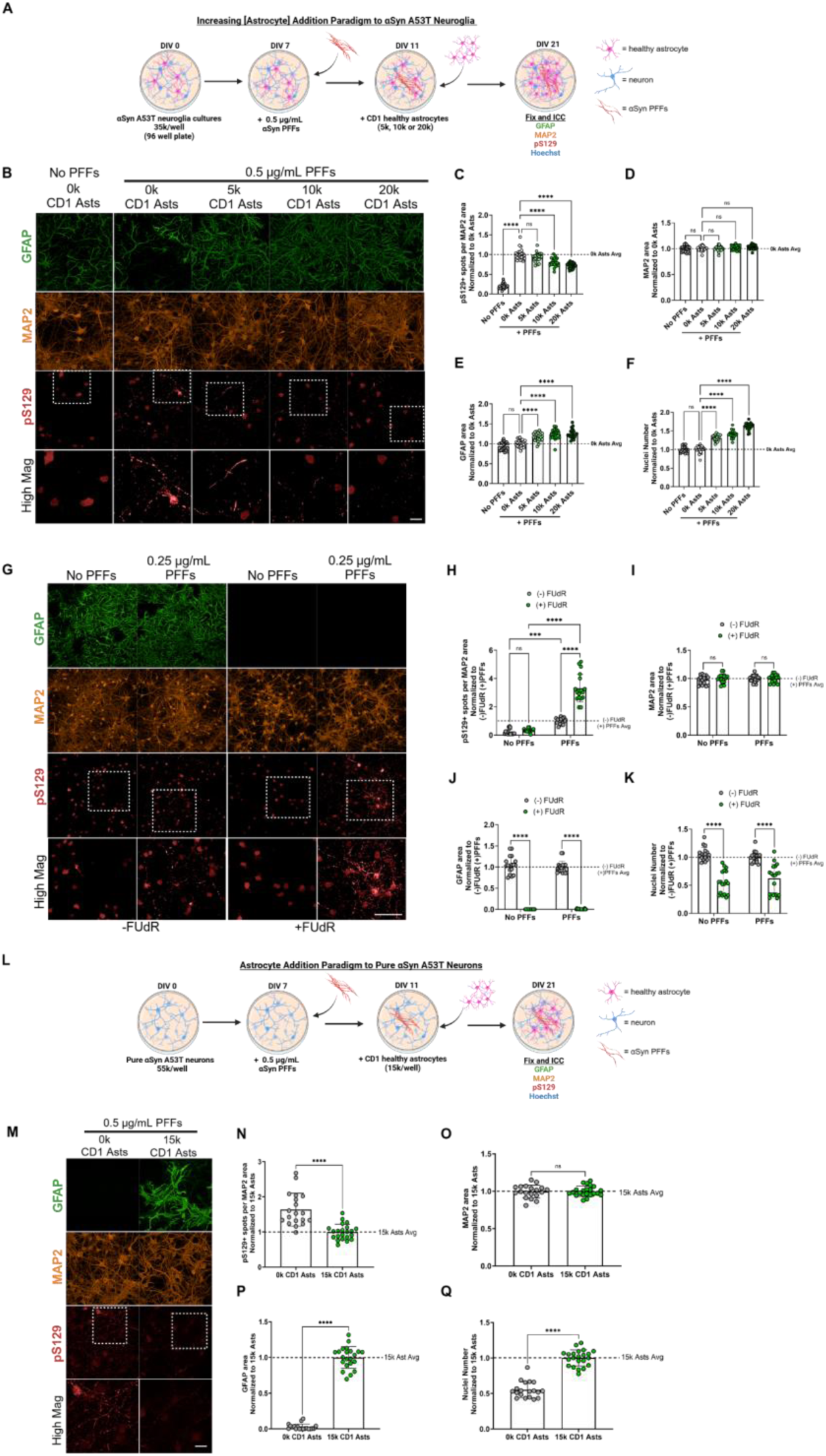
Primary astrocytes prevent pS129 pathology in neurons induced by αSyn PFFs. A) Schematic of the assay used in B-F. B) Representative images of DIV 21 neuroglia cultures with increasing densities of primary astrocytes (Asts) added and immunocytochemically stained for pS129, GFAP, MAP2, and nuclei with Hoechst. Scale bar = 50 µm. Quantification pS129+ spots per MAP2 area (C), MAP2 area (D), GFAP area (E), and nuclei number (F), expressed as a fold change of the 0k Ast group. N = 3-6 wells/plate across 3 separate experiments. One-way ANOVA with Sidak post hoc, ****= p < 0.0001. G) Representative images of DIV 21 neuroglia cultures (-FUdR) and pure neuron cultures (+ FUdR) and immunocytochemically stained for pS129, GFAP, MAP2, and nuclei with Hoechst. Scale bar = 50 µm. Quantification pS129+ spots per MAP2 area (H), MAP2 area (I), GFAP area (J), and nuclei number (K), expressed as a fold change of the (-) FUdR) / (+) PFF group. N = 3-6 wells/plate across 3 separate experiments. Two-way ANOVA with Sidak post hoc, ***= p < 0.001, ****= p < 0.0001. L) Schematic of the assay used in M-Q. M) Representative images of DIV 21 pure neuron cultures -/+ Asts and immunocytochemically stained for pS129, GFAP, MAP2, and nuclei with Hoechst. Scale bar = 50 µm. Quantification pS129+ spots per MAP2 area (N), MAP2 area (O), GFAP area (P), and nuclei number (Q), expressed as a fold change of the 15k Asts group. N = 3-10 wells/plate across 3 separate experiments. Unpaired student’s t-test, ****= p < 0.0001.

Next, we hypothesized that depletion of A53T αSyn transgenic astrocytes from neuroglia cultures would enhance endogenous αSyn pS129 pathology in primary neurons, subsequently increasing the efficacy of exogenous astrocytes when added. To test this, we used the mitotic inhibitor 5-Fluoro-2’deoxyuridine (FUdR) to deplete proliferating glia cells prior to plating neurons. Endogenous αSyn pS129+ pathology was examined in FUdR-treated pure neuron cultures compared to untreated cultures induced by αSyn PFFs. Representative images and quantification show more pS129+ spots in FUdR-treated neuron cultures compared to untreated cultures (Figure 3G,H). Therefore, depletion of A53T αSyn transgenic astrocytes increased endogenous αSyn pS129+ pathology in neurons.

Next, we examined whether addition of astrocytes to pure neurons modulated endogenous αSyn pS129+ pathology to a greater extent than in the neuroglia cultures after adding additional astrocytes. Following the paradigm depicted in figure 3L, astrocytes were added four days after pure neurons were treated with αSyn PFFs to allow pure neurons to take up most of the αSyn PFFs prior to astrocyte addition; Cells were subsequently immunostained for GFAP, MAP2, or pS129 on DIV 21. Representative images and quantification of cultures with astrocytes added show fewer pS129+ spots compared to pure neuron control (Figure 3M,N). Astrocyte addition to pure neuron cultures more robustly decreased neuronal αSyn pS129+ pathology (by ∼50%) compared to astrocyte addition to neuronal cultures with A53T αSyn transgenic astrocytes still present. This data confirmed our hypothesis that astrocytes can reduce neuronal exposure to propagating αSyn seeds.

### LC astrocytes do not mitigate pS129 pathology in neurons induced by αSyn PFFs

We next hypothesized that LC astrocytes would be unable to prevent endogenous αSyn pS129+ pathology in neurons. We tested this by treating astrocytes with CQ or CA-074 overnight prior to adding to neuroglia cultures already exposed to PFFs (Figure 4A). Representative images and quantification of neuroglia cultures treated with LC astrocytes had more pS129+ spots compared to neuroglia cultures with unimpaired (vehicle-treated) astrocytes added (Figure 4B,C). This suggests that lysosomal function plays a critical role in the ability of astrocytes to reduce endogenous αSyn seeding in neurons.

**Figure 4.**
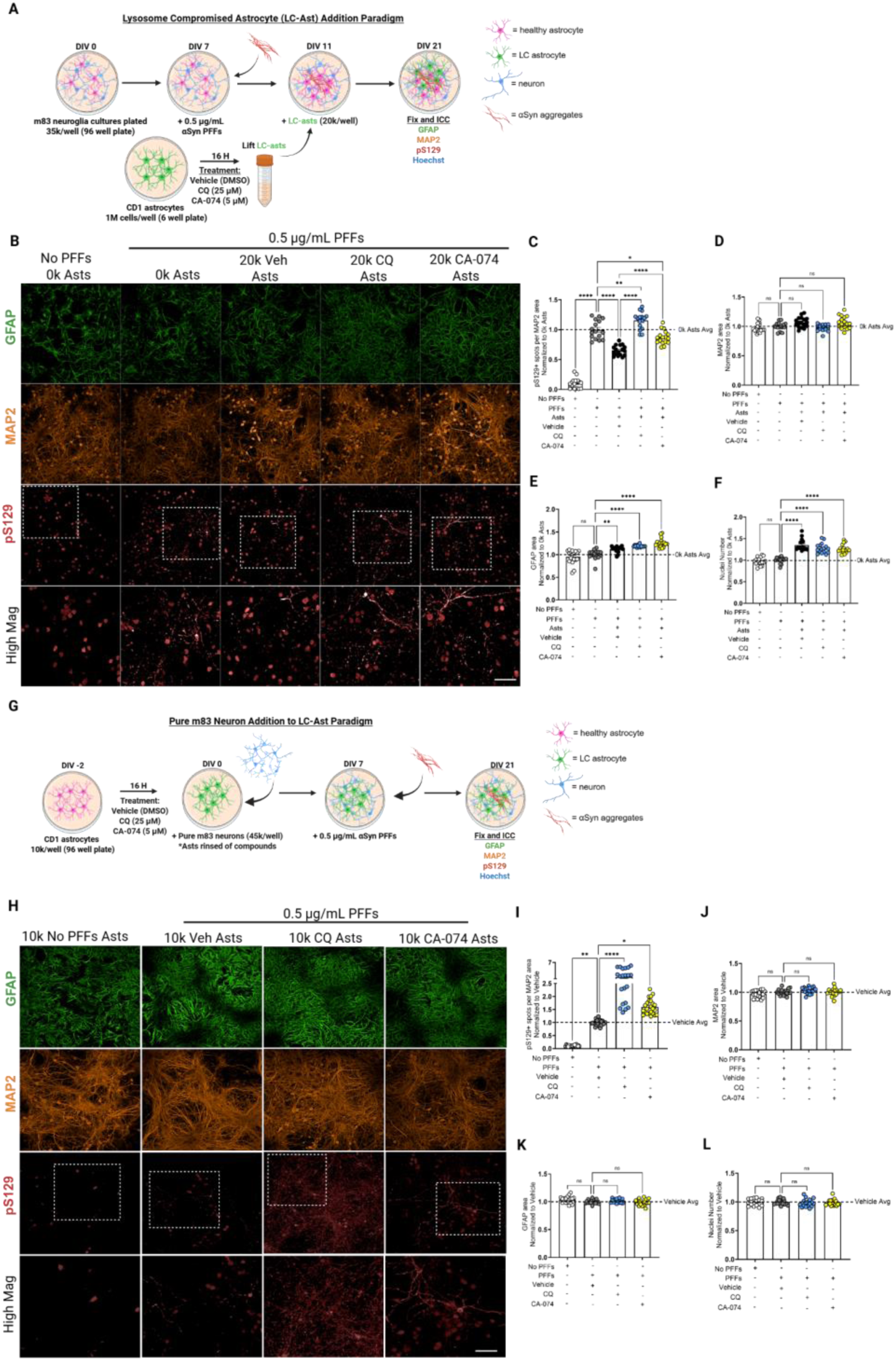
LC astrocytes do not prevent pS129 pathology in neurons induced by αSyn PFFs. A) Schematic of the assay used in B-F. B) Representative images of DIV 21 neuroglia cultures -/+ healthy or LC Asts and immunocytochemically stained for pS129, GFAP, MAP2, and nuclei with Hoechst. Scale bar = 50 µm. Quantification pS129+ spots per MAP2 area (C), MAP2 area (D), GFAP area (E), and nuclei number (F), expressed as a fold change of the 0k Ast group. N = 3-6 wells/plate across 3 separate experiments. One-way ANOVA with Sidak post hoc, *= p < 0.05, **= p < 0.01, ****= p < 0.0001. G) Schematic of the assay used in H-L. H) Representative images of DIV 21 neuron cultures + healthy or LC Asts and immunocytochemically stained for pS129, GFAP, MAP2, and nuclei with Hoechst. Scale bar = 50 µm. Quantification pS129+ spots per MAP2 area (I), MAP2 area (J), GFAP area (K), and nuclei number (L), expressed as a fold change of the vehicle-treated Ast group. N = 3-6 wells/plate across 4 separate experiments. One-way ANOVA with Dunnett’s post hoc, *= p < 0.05, **= p < 0.01, ****= p < 0.0001.

Coming from cultures with a mixed impaired and unimpaired astrocyte population, we next used a model to assess how solely LC astrocytes would affect endogenous αSyn pS129+ pathology in neurons. In this paradigm, astrocytes were plated first and treated overnight with CQ or CA-074, rinsed and pure neurons were plated on top. Co-cultures were exposed to αSyn PFFs and subsequently immunostained for GFAP, MAP2, or pS129 (Figure 4G). Representative images and quantification of co-cultures with LC astrocytes show increased pS129+ spots compared to co-cultures with vehicle-treated astrocytes (Figure 4H,I). Therefore, astrocytes with insufficient lysosome capacity have a reduced proficiency to mitigate αSyn pathology in neuronal cultures as well as astrocytes with normal lysosome function.

### αSyn PFF preloaded LC astrocytes induce endogenous αSyn pS129 pathology in neurons

Since LC astrocytes accumulate αSyn PFFs following exposure to PFFs (Figure 2) and lose the ability to alleviate pS129 pathology in neurons (Figure 4), we examined whether LC astrocytes preloaded with αSyn PFFs could spread seed-competent αSyn to neurons. In this assay, astrocytes are loaded with αSyn PFFs, rinsed, and treated overnight with CQ. αSyn PFF preloaded astrocytes are added to neuroglia cultures and immunostained on for GFAP, MAP2, or pS129 (Figure 5A). Quantification reveals the presence of pS129+ spots only in neuroglia cultures with PFF-preloaded, CQ-treated astrocytes compared to PFF-preloaded, vehicle-treated astrocytes, highlighting the importance of a functional lysosome to neutralize aSyn aggregates (Figure 5B,C).

**Figure 5.**
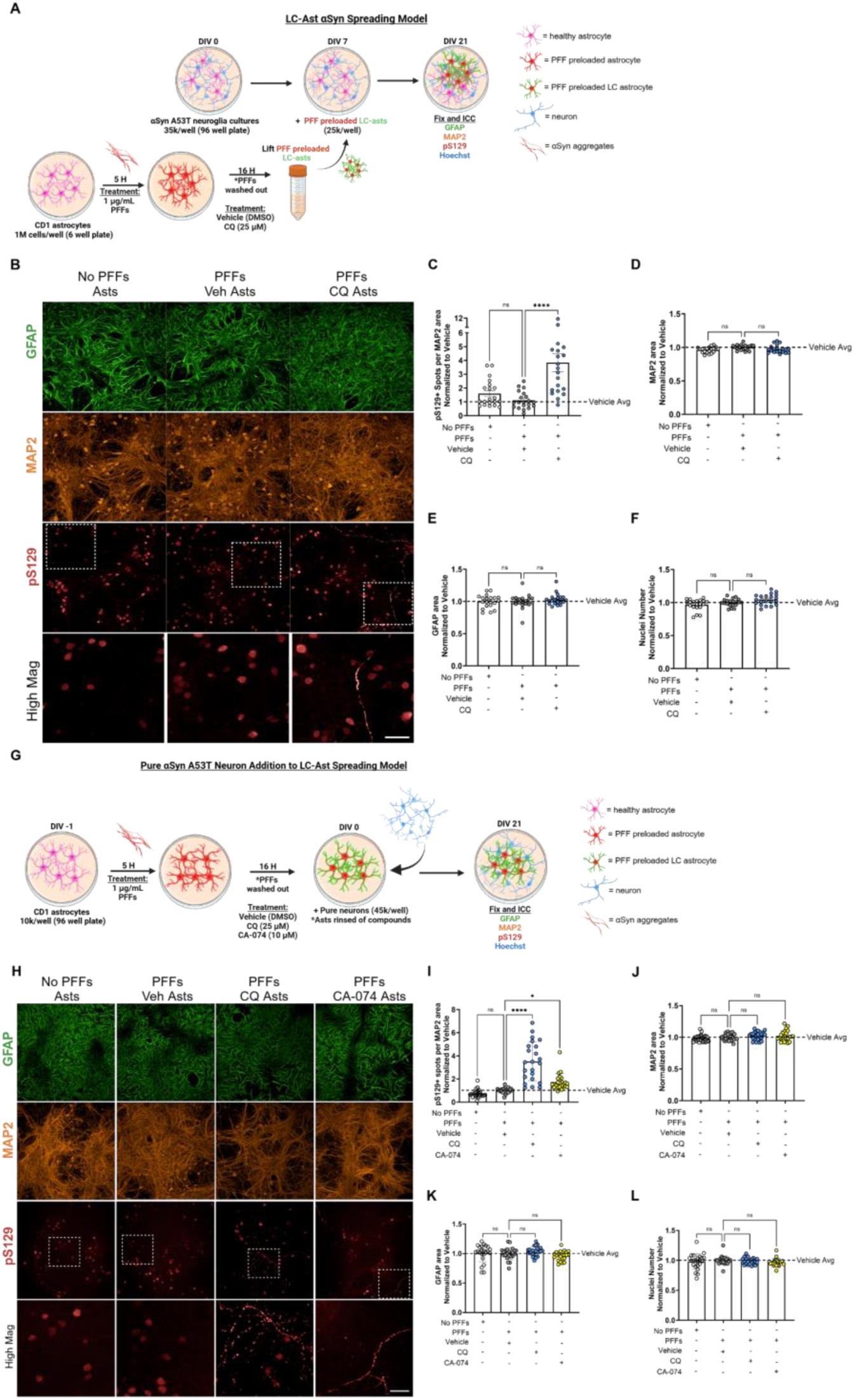
αSyn PFF preloaded LC astrocytes induce pS129 pathology in neurons. A) Schematic of the assay used in B-F. B) Representative images of DIV 21 neuroglia cultures + αSyn PFF-preloaded Asts and immunocytochemically stained for pS129, GFAP, MAP2, and nuclei with Hoechst. Scale bar = 50 µm. Quantification pS129+ spots per MAP2 area (C), MAP2 area (D), GFAP area (E), and nuclei number (F), expressed as a fold change of the vehicle-treated Ast group. N = 3-6 wells/plate across 3 separate experiments. One-way ANOVA with Dunnett’s post hoc, ****= p < 0.0001. G) Schematic of the assay used in H-L. H) Representative images of DIV 21 neuron cultures + αSyn PFF-preloaded Asts and immunocytochemically stained for pS129, GFAP, MAP2, and nuclei with Hoechst. Scale bar = 50 µm. Quantification pS129+ spots per MAP2 area (I), MAP2 area (J), GFAP area (K), and nuclei number (L), expressed as a fold change of the vehicle-treated Ast group. N = 3-6 wells/plate across 4 separate experiments. One-way ANOVA with Dunnett’s post hoc, *= p < 0.05, ****= p < 0.0001.

Finally, since the paradigm in figure 5A includes the presence of A53T αSyn transgenic astrocytes, we decided to use the same assay depicted in figure 4G-L to better assess the ability of solely LC astrocytes to induce endogenous αSyn pS129 pathology in pure neurons. In this experiment, we plated astrocytes first, preloaded them with αSyn PFFs, rinsed, and treated with CQ or CA-074 overnight. Pure neurons were added on top of the astrocytes and the cells were immunostained for GFAP, MAP2, or pS129 (Figure 5G). Similarly, only neurons cultured with PFF pre-loaded astrocytes treated with CQ and to a lesser degree with CA-074 demonstrated αSyn pS129 pathology (Figure 5H-I), further suggesting that LC astrocytes can spread seed-competent αSyn to neurons. Taken together, these data suggest that astrocytic lysosome efficiency is important for degrading αSyn aggregates and slowing the spread of αSyn pathology in neurons.

## Discussion

Using co-culture models to specifically manipulate astrocytes, we provide evidence that astrocytic lysosomal deficiency causes the accumulation of αSyn PFFs, which can lead to the spread of seed-competent αSyn to surrounding neurons for formation of endogenous pS129+ aggregation. We show that healthy WT mouse and human astrocytes can readily take up and preferentially degrade αSyn PFFs *in vitro* and exogenously added WT astrocytes can mitigate PFF-induced aggregation of endogenous αSyn in primary neurons. However, either general astrocytic lysosomal damage and dysfunction or specific minor lysosome enzymatic inhibition disrupts αSyn PFF degradation, resulting in an elevation of endogenous αSyn aggregation in neurons and spread of αSyn pathology. This work emphasizes the importance of the astrocytic lysosomal pathway in the clearance αSyn aggregates and supports a non-cell autonomous contribution to PD neuropathology.

Firstly, we observed that astrocytes take up more αSyn PFFs over time than αSyn monomer (Figure 1A-C). Due to the use of the pHrodo tag, we know that both forms of αSyn were taken up through acidic compartments including the endosome, which agrees with previous reports of αSyn uptake mechanisms (Lee *et al*, 2008b, 2008a; Andromidas *et al*, 2024; Koob & Sacchetti, 2018). Interestingly, the observations that monomeric αSyn is taken up less than aggregates and that it is not degraded in the same manner suggest that the two αSyn formats are differentially internalized and processed (Figure1D-F). Also, in αSyn PFF-treated astrocytes, we observed that total αSyn staining localized to LAMP1+ compartments suggesting that αSyn PFFs are targeted to lysosomes for degradation (Figure 1G-H). Taken together, astrocytes specifically and preferentially target αSyn PFFs, and not monomeric αSyn, to the lysosome and degrade them. This specific difference in the astrocytic metabolism of αSyn aggregates led us to explore the role of lysosome degradation further.

Due to the strong genetic and mechanistic evidence for Cat D and Cat B involvement in PD, (Lin *et al*, 2024; Nalls *et al*, 2019; Bunk *et al*, 2021; Jones-Tabah *et al*, 2024; McGlinchey & Lee, 2015; Drobny *et al*, 2022; McGlinchey *et al*, 2019) inhibitors of Cat B (Leu or CA-074) and an inhibitor of Cat D (Pep A) were utilized to understand cathepsin-specific enzymatic digestion of αSyn aggregates in astrocytes. The lysosomotropic compound CQ was used to examine how general lysosome deficiency affects astrocytic αSyn PFF degradation, as it is commonly used to raise lysosomal pH to limit enzymatic digestion (Zeng *et al*, 2020). In αSyn PFF and CQ-, Leu- or CA-074-treated astrocytes, we observed unique partially degraded forms of αSyn not detected in αSyn PFF and vehicle- or Pep A-treated astrocytes (Figure 2). Based on the antibodies used in this study (Extended View Figure 1A), these degraded forms of αSyn are likely c-terminally fragmented since they were not seen when αSyn PFF-treated astrocyte lysates were probed with an αSyn antibody targeting a c-terminal epitope (MJFR1 epitope aa118-123; Extended View Figure 1B-C). Intriguingly, this accumulation of unique degraded forms of αSyn was only observed with Cat B inhibition (Leu or CA-074 treatment) and not with Cat D inhibition (Pep A treatment) (Figure 2). Despite the compelling evidence for the degradation of αSyn aggregates by Cat D (Bunk *et al*, 2021; Qiao *et al*, 2008; Cullen *et al*, 2009), it’s possible that Cat B plays a larger role in the context of astrocytic αSyn degradation. Moreover, other cathepsins could play a role in astrocytic αSyn degradation and compensate for Cat B (Matsuki *et al*, 2024). A full analysis of cathepsin activity in astrocytes regarding αSyn digestion may be warranted.

A significant distinction in our study is that we did not observe overt changes to lysosome morphology or damage with αSyn PFF treatment alone (Extended View Figure 2). Many studies of glia cells exposed to various forms of αSyn observed organellar dysfunction or alterations in the secretome (Lindström *et al*, 2017; Freeman *et al*, 2013; Raj *et al*, 2024; Chavarría *et al*, 2018). These studies all used different PFFs that were exposed to glia cells at much higher concentrations than this study, which may account for the differing results. However, this work adds to a growing number of studies that suggest glia cell-specific lysosomal dysfunction plays a critical function in the contribution to neurodegenerative disease pathology and progression (Quick *et al*, 2023; Uddin & Lim, 2022; Balta & Zunke, 2021; Aflaki *et al*, 2020).

Importantly, only LC astrocytes preloaded with αSyn PFFs induced spread of αSyn pathology (Figure 5). This observation is aligned with our data in figure 2, clearly showing that LC astrocytes accumulated αSyn at times where the vehicle-treated astrocytes degraded αSyn PFFs rapidly and efficiently. It has been suggested that truncated forms of αSyn are more prone to aggregation (Sorrentino & Giasson, 2020; Liu *et al*, 2005; Zhang *et al*, 2022). Moreover, Cat B-digested αSyn PFFs form truncated αSyn that has been shown to increase aggregation propensity in neurons (Tsujimura *et al*, 2015). It is possible that these unique partially degraded forms of αSyn PFFs, only observed in LC astrocytes, play a significant role in the spread of αSyn pathology in our model. Furthermore, αSyn PFF preloaded CQ-treated astrocytes induced more pS129 αSyn pathology in neurons compared to αSyn PFF preloaded CA-074-treated astrocytes (Figure 5G-L). Since CQ treatment caused overt changes to lysosome size, number, and lysosome damage (as demonstrated with Gal3 localization) that were absent following CatB inhibition, it is possible that the mechanism by which these astrocytes induced αSyn pathology in neurons is different, even though both compounds clearly led to accumulation of non-degraded and degraded αSyn aggregates (Figure 2). A proposed model for propagation based on our findings is shown in the synopsis image. However, the potential mechanisms of αSyn aggregate spread remain to be elucidated.

In PD, αSyn aggregates are hypothesized to spread in a prion-like manner through synaptically coupled networks of neurons leading to LB and LN pathology and neurodegeneration (Henrich *et al*, 2020). However, it’s suggested that there is no clear correlation between the strength of the synaptic connections and the magnitude or persistence of the propagated pathology, which implies that spread of αSyn is not solely through neuronal synaptic connectivity (Henrich *et al*, 2020). Additionally, there is evidence suggesting that αSyn is transmitted amongst neurons, astrocytes, and microglia, but the mechanisms by which this occurs remain intensely debated (Dehay *et al*, 2015; Loria *et al*, 2017; Rostami *et al*, 2017; Walsh & Selkoe, 2016; Emmanouilidou *et al*, 2010; El-Agnaf *et al*, 2003). Many of these studies report αSyn is released via exosomes, by lysosome exocytosis following lysosome dysfunction or through neuroinflammation (Burbidge *et al*, 2022; Alvarez-Erviti *et al*, 2011; Xie *et al*, 2022; Bae *et al*, 2022; Lee *et al*, 2014). Moreover, two other studies show that neuron to neuron or astrocyte to astrocyte αSyn propagation can occur through direct contact with tunneling nanotubes (Abounit *et al*, 2016; Rostami *et al*, 2017). In future studies, we aim to further investigate these possible mechanisms of astrocytic αSyn PFF spread in our model.

This work supports a growing body of research suggesting a role for glia cells, specifically astrocytes, in PD etiology and disease progression. We illustrate that overt lysosome damage or Cat B inhibition in astrocytes can modulate and spread αSyn pathology in neurons. Astrocytes robustly take up and degrade αSyn aggregates via the endolysosomal system and that enhancement of this pathway can be harnessed to develop new disease-modifying therapeutics for PD.

## Supporting information

Extended View Figure 1

Extended View Figure 2

Synopsis Image

## Author contributions

L.M.R., O.M, J.S, and J.S* contributed to the study conception and design. Material preparation, data collection, and analysis were performed by L.M.R. and O.M. The first draft of the manuscript was written by L.M.R. and J.S*. All authors commented on previous versions of the manuscript. All authors read and approved of the final manuscript.

## Acknowledgements.

Jordan Dasilva in the Comparative Medicine East group at AbbVie for her help with the isolation of primary neuroglia cultures. BioRender was used to create all assay schematics in the figures and the synopsis image.

