## Supplementary figures and images for "Astrocytic lysosome deficits reduce alpha-synuclein degradation and induce spread of pathology"

### Extended View Figure 1

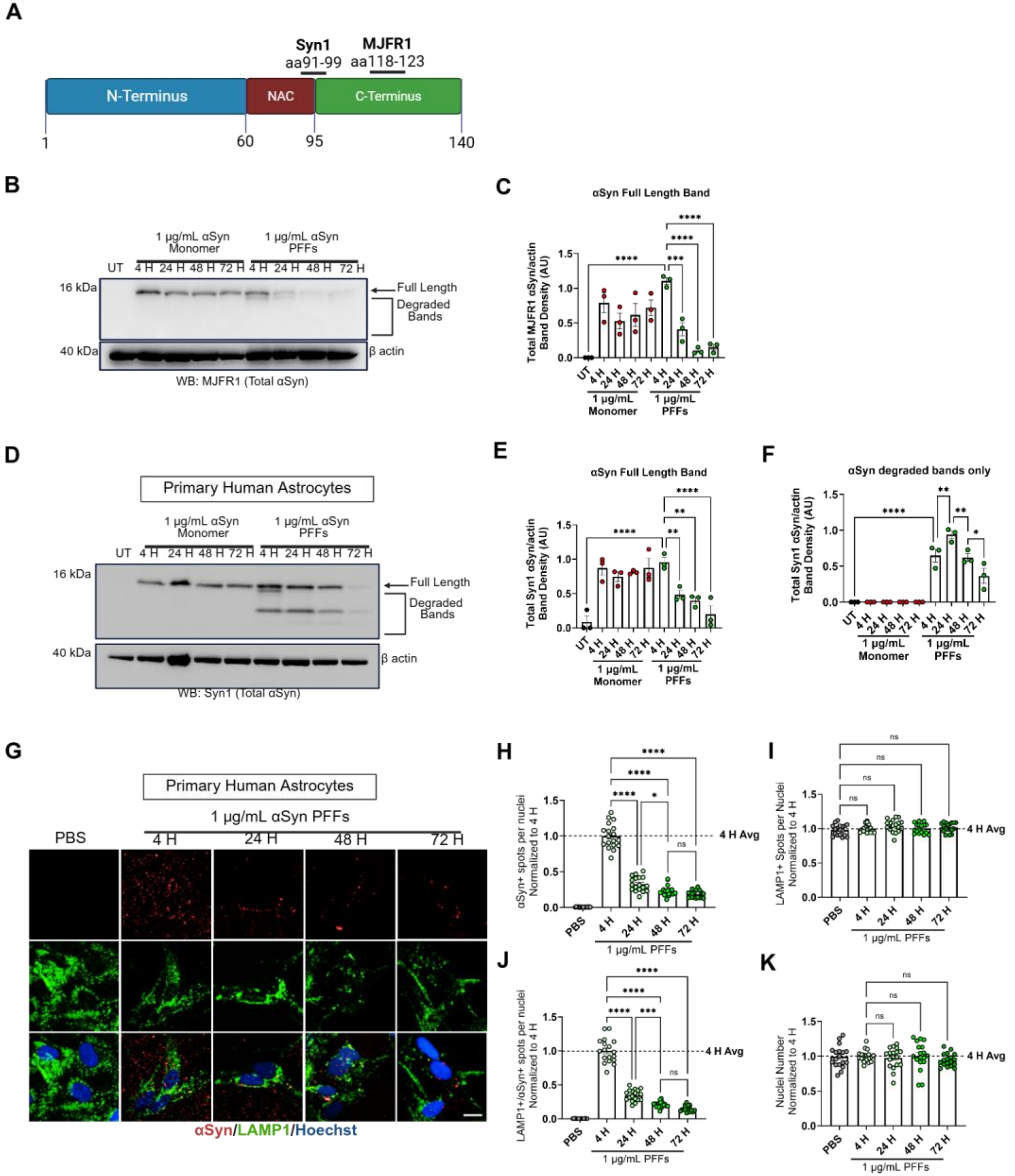

### Extended View Figure 2

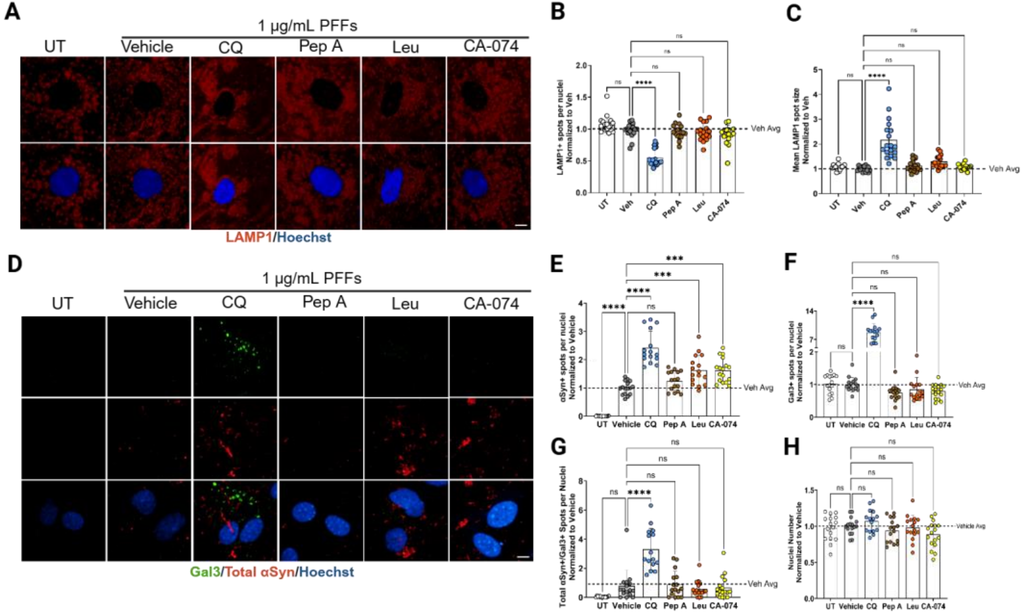

### Synopsis Image

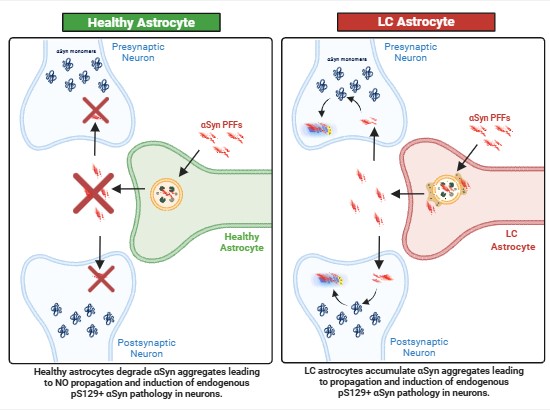
